# A rare variant in ANP32B impairs influenza virus replication in human cells

**DOI:** 10.1101/2020.04.06.027482

**Authors:** Ecco Staller, Laury Baillon, Rebecca Frise, Thomas P. Peacock, Carol M. Sheppard, Vanessa Sancho-Shimizu, Wendy Barclay

**Affiliations:** Department of Infectious Disease, Faculty of Medicine, Imperial College London, United Kingdom

## Abstract

Viruses require host factors to support their replication, and genetic variation in such factors can affect susceptibility to infectious disease. Influenza virus replication in human cells relies on ANP32 proteins, which are involved in assembly of replication-competent dimeric influenza virus polymerase (FluPol) complexes. Here, we investigate naturally occurring single nucleotide variants (SNV) in the human *Anp32A* and *Anp32B* genes. We note that variant rs182096718 in *Anp32B* is found at a higher frequency than other variants in either gene. This SNV results in a D130A substitution in ANP32B, which is less able to support FluPol activity than wildtype ANP32B and binds FluPol with lower affinity. Interestingly, ANP32B-D130A exerts a dominant negative effect over wildtype ANP32B and interferes with the functionally redundant paralogue ANP32A. FluPol activity and virus replication are attenuated in CRISPR-edited cells expressing wildtype ANP32A and mutant ANP32B-D130A. We propose a model in which the D130A mutation impairs FluPol dimer formation, thus resulting in compromised replication. We suggest that both homozygous and heterozygous carriers of rs182096718 may have some genetic protection against influenza viruses.

## INTRODUCTION

All viruses rely on host factors to support their replication. Genetic variation in such proteins can affect susceptibility to infectious disease (; Ciancanelli, Abel, Zhang, & Casanova, 2016; Karlsson, Kwiatkowski, & Sabeti, 2014; Pittman, Glover, Wang, & Ko, 2016; Q. Zhang, 2020). Perhaps the best-known mutation conferring genetic resistance to viral infection is the Δ32 variant of the HIV-1 co-receptor C-C chemokine receptor type 5 (*CCR5*) (Carrington, Dean, Martin, & O’Brien, 1999). Single nucleotide variants (SNV) that reduce susceptibility to virus infection include a P424A substitution in the filovirus endosomal fusion receptor Niemann-PickC1 (*NPC1*) (Carette et al., 2011; Kondoh et al., 2018), and a G428A mutation in fucosyltransferase 2 (*FUT2*) that renders homozygous carriers resistant to norovirus (Lindesmith et al., 2003; Thorven et al., 2005). These variants affect viral entry – no SNVs resulting in defects in replication have thus far been described.

Genetic variation in the endosomal transmembrane protein IFITM3 has been associated with increased severity of infection with influenza virus. The single nucleotide polymorphism (SNP) rs12252 leads to an N-terminal 21-amino acid truncation through an alteration of the first splice acceptor site, and is perhaps the best known genetic variant affecting influenza disease severity (Everitt et al., 2012; Lee et al., 2017; Y.-H. Zhang et al., 2013). Although the association has not always been clear (Mills et al., 2014; Randolph et al., 2017), a recent meta-analysis including 12 studies and over 16,000 subjects confirmed the link between the IFITM3 minority allele and influenza severity (Prabhu, Chakraborty, Kumar, & Banerjee, 2018). Another important polymorphism in the IFITM3 gene associated with severe influenza is located in the 5’ UTR and leads to transcriptional repression through increased CTCF binding to the promoter (Allen et al., 2017).

Some non-synonymous natural variants in the restriction factor MxA lost antiviral activity, and even exerted a dominant-negative effect on the antiviral activity of wildtype MxA, suggesting heterozygous carriers of these variants may be more susceptible to influenza infection (Graf et al., 2018). Additional mutations associated with severe influenza (reviewed in (Ciancanelli et al., 2016; Q. Zhang, 2020)) have been described in the complement factor CD55/DAF (Chatzopoulou, Gioula, Kioumis, Chatzidimitriou, & Exindari, 2019), the pulmonary surfactant protein SFTPA2 (Herrera-Ramos et al., 2014), the protease TMPRSS2 (Cheng et al., 2015), the endothelial transcriptional activator GATA2 (Sologuren et al., 2018), interferon regulators IRF7 and IRF9 (Bravo Garcia-Morato et al., 2019; Ciancanelli et al., 2015; Hernandez et al., 2018), and the endosomal dsRNA receptor TLR3 (Esposito et al., 2012; Hidaka et al., 2006; Lim et al., 2019).

Almost all the host genetic variation relevant to influenza that has been described is associated with exacerbating rather than alleviating influenza virus infection, a bias explained by the higher visibility of severe cases, and the opportunity to perform whole genome sequencing on patients with severe symptoms. Hence, besides a pair of polymorphisms in the non-coding region (NCR) of the Galectin-1 encoding gene *LGALS1*, which may confer some protection against H7N9 influenza A virus (IAV) in Chinese poultry workers (Y. Chen et al., 2015; Nogales & M, 2019), no single nucleotide variants protective against influenza virus have thus far been described.

Influenza replication and transcription are carried out in the host cell nucleus by the virus-encoded heterotrimeric RNA-dependent RNA polymerase (FluPol) (reviewed in (Fodor & Te Velthuis, 2019; Wandzik, Kouba, & Cusack, 2020), which relies on host factors to support RNA synthesis (Staller and Barclay, 2020, *in press*) (Peacock, Sheppard, Staller, & Barclay, 2019). Oligomerisation of trimeric FluPol molecules into dimers, trimers or even higher order complexes is now believed to be required for efficient replication (Carrique et al, 2020, *in press*) (Chang et al., 2015; K. Y. Chen, Santos Afonso, Enouf, Isel, & Naffakh, 2019; Fan et al., 2019; Peng et al., 2019).

Influenza viruses co-opt the host proteins acidic nuclear phosphoprotein 32 kilodaltons A (ANP32A) and ANP32B to support their replication (J. S. Long et al., 2016; Staller et al., 2019; H. Zhang et al., 2019). ANP32A and ANP32B are functionally redundant in their support for influenza virus: in human cells lacking either but not both ANP32A (AKO) or ANP32B (BKO), virus proliferation is not impaired. In cells lacking both proteins (dKO), however, FluPol activity and virus replication are completely abrogated (Staller et al., 2019; H. Zhang et al., 2019). ANP32 proteins are expressed in many different human tissues, including fibroblasts and lung (Lonsdale et al., 2013), and involved in a wide range of cellular processes, including apoptosis and transcriptional regulation (reviewed in (Reilly, Yu, Hamiche, & Wang, 2014)). All of the human paralogues – ANP32A, ANP32B and ANP32E – associate with histones (Kleiner, Hang, Molloy, Chait, & Kapoor, 2018; Obri et al., 2014; Saavedra et al., 2017; Tochio et al., 2010), placing them at the chromatin within the nucleus.

ANP32A and ANP32B are 249 and 251 amino acids long, respectively, with two major domains: a structured N-terminal leucine-rich repeat region (LRR) and an intrinsically disordered low-complexity acidic region (LCAR) (Camacho-Zarco et al., 2020; Huyton & Wolberger, 2007; Tochio et al., 2010). Mutational analysis of ANP32 proteins of several species has shown that the identity of amino acid residues 129 and 130 dictates pro-viral function (Jason S. Long et al., 2019; Thomas P. Peacock et al., 2020; Staller et al., 2019; H. Zhang et al., 2019). ANP32 proteins that cannot be co-opted by influenza virus include human ANP32E, chicken ANP32B, and mouse ANP32A. What these orthologues have in common is divergence from the pro-viral dyad 129N-130D: human ANP32E has glutamate at position 129 (129E) rather than asparagine, chicken ANP32B has isoleucine at position 129 (129I) and asparagine at position 130 (130N), and mouse ANP32A has alanine at position 130 (130A), suggesting that these amino acids are important for pro-viral activity of ANP32 proteins.

Recent structural work depicts the ANP32A LRR domain as stabilising a replication-competent dimer of a viral promoter-bound ‘replicating’ FluPol heterotrimer (FluPol^R^) and an ‘encapsidating’ FluPol^E^, which is proposed to receive the nascent viral RNA product synthesised by FluPol^R^ (Carrique et al, 2020, *in press*). More specifically, the N-terminal half of the LRR interacts with FluPol^R^, while the C-terminal portion interacts with FluPol^E^, in effect bridging the dimer. The importance of residues 129 and 130 is explained by direct interaction with basic residues on the PB1 subunit of FluPol^E^. Interaction of the LCAR domain of ANP32A with FluPol has also been postulated (Camacho-Zarco et al., 2020; Mistry et al., 2020) (Carrique et al, 2020, *in press*).

Here we describe natural variation in the human *Anp32A* and *Anp32B* genes, demonstrating that the naturally occurring SNV rs182096718 in *Anp32B* can confer protection from influenza virus replication by compromising the interaction between FluPol and the host factor. We demonstrate that rs182096718 exerts a dominant-negative effect over wildtype ANP32B and moreover interferes with the functionally redundant paralogue ANP32A through paralogue interference (Kaltenegger & Ober, 2015). We propose a model explaining why influenza replication is impaired in cells expressing the mutant ANP32B protein.

## RESULTS

### A relatively common single nucleotide variant encodes ANP32B-D130A

We searched public databases for rare missense SNVs in the human *Anp32A* and *Anp32B* genes that were predicted, on the basis of our previous work, to have functional impact on the pro-influenza virus activity of the proteins. We predominantly used the genome aggregation database gnomAD v2, which holds 125,748 whole exomes and 15,708 whole genomes from unrelated individuals sequenced in disease-specific or population genetic studies (Karczewski et al., 2019). The gnomAD database identifies 54 missense SNVs in *Anp32A* and 82 in *Anp32B* (observed/expected score of 0.39 vs 0.68), suggesting that the former may be more conserved. It is not currently possible to obtain specific phenotypic information about individual carriers of a particular SNV from gnomAD.

Four SNVs in *Anp32A* and five in *Anp32B* were selected for investigation, taking into account changes in charge, bulk, polarity and putative solvent exposure of the encoded amino acid substitutions (Huyton & Wolberger, 2007; Tochio et al., 2010), as well as the previously established importance of positions 129 and 130 for pro-influenza viral activity (Figure 1A). All the variants in *Anp32A* are exceedingly rare (MAF<0.001), and no homozygous carriers have been identified. The serine mutants at position 158, S158A and S158T were of interest as this is a putative phosphorylation site (Hong et al., 2004). The SNV responsible for a mutant ANP32A-D130N protein (rs771143708) is found in a single heterozygous individual out of 3,854 in the Avon Longitudinal Study of Parents and Children (ALSPAC) cohort. This SNV is not described in the much larger gnomAD database, although it is cited in the NCBI database dbSNP. rs751239680 (ANP32A-R132Q) is found in a single heterozygous individual out of 10,052 in the African cohort of the gnomAD database, rs772530468 (ANP32A-S158T) is present in a single South Asian male in 23,070 in the gnomAD database, as well as two separate homozygous individuals in the Trans-omics for precision medicine (TOPMed) database (Taliun et al., 2019). Finally, a single African male in 5,032 in the gnomAD database is a heterozygous carrier of rs772530468 (ANP32A-S158A) (Figure 1B).

**Figure 1.**
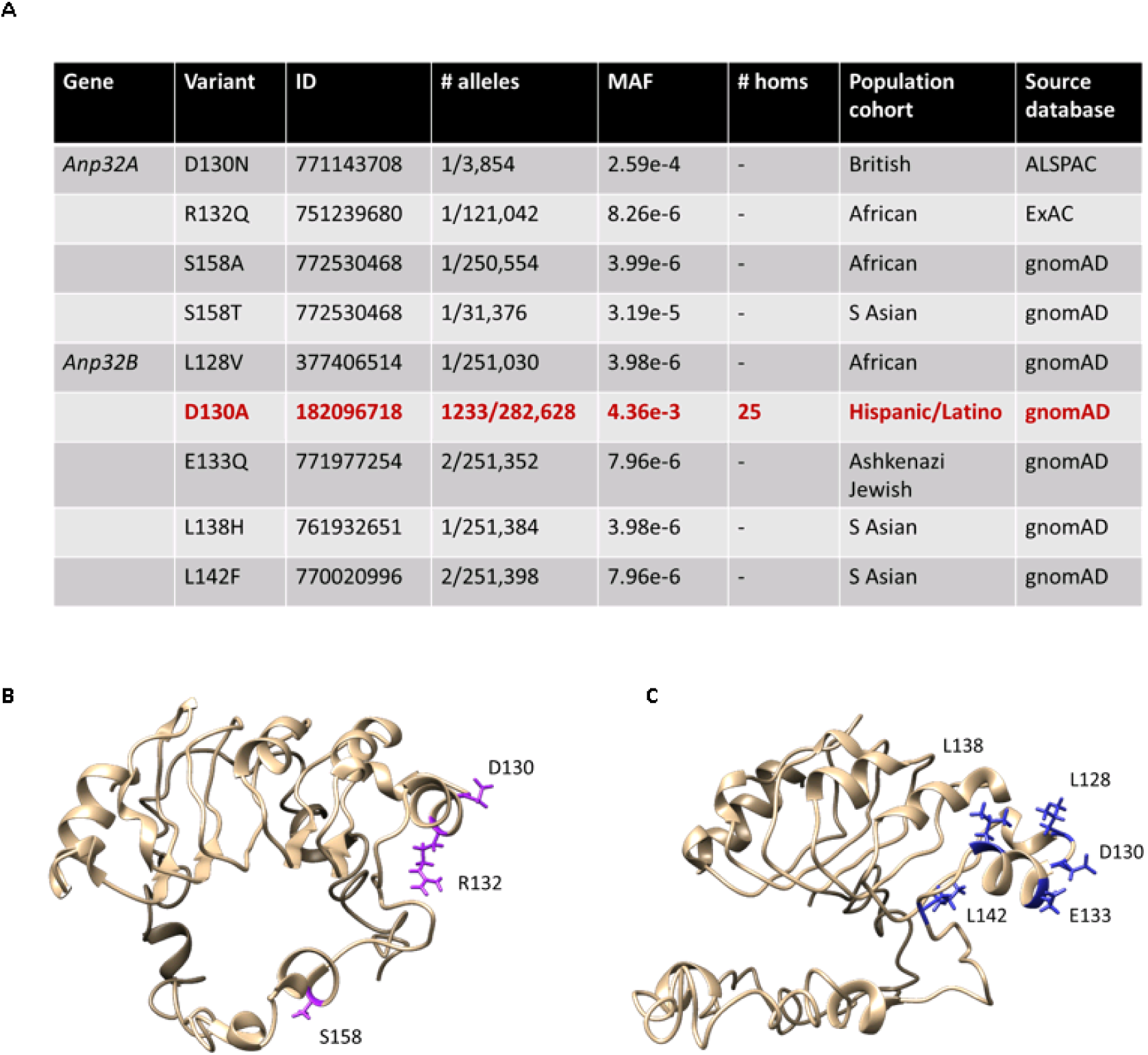
Selected naturally occurring missense single nucleotide variants in *Anp32A* and *Anp32B*. (A) Amino acid substitutions (variant), NCBI dbSNP Reference IDs, number of minority alleles, global minor allele frequency (MAF), number of homozygous carriers (homs), predominant gnomAD population cohort in which the variant is found, and the main source database for each variant (B) Structural model of ANP32A showing amino acids affected by selected SNVs (C) Structural model of ANP32B showing amino acids affected by selected SNVs

Most variants in *Anp32B* were also low-frequency (MAF<0.0001): rs377406514 (ANP32B-L128V) is found in a single African female out of 10,066 in the gnomAD database; rs771977254 (ANP32B-E133Q) occurs in two out of 4,900 Ashkenazi Jewish females in the gnomAD database; rs761932651 (ANP32B-L138H) is found in one South Asian male in 23,070 in the gnomAD database; rs770020996 (ANP32B-L142F) occurs in one female in 7,544 and one male in 23,072 from the South Asian cohort of the gnomAD database (Figure 1C).

There was, however, an interesting exception. SNV rs182096718, encoding ANP32B-D130A, was relatively common in the Hispanic / Latino cohort of the gnomAD database, where in a total pool of 35,420 individuals, 1,209 heterozygous carriers were identified, as well as 25 homozygous carriers. The D130A substitution in ANP32B is thus present in 3.41% of the Latino cohort. The global MAF of this variant is 0.0044 which is much higher than any other SNV in either *Anp32A* or *Anp32B*.

### Natural variants of ANP32 proteins affect support of FluPol activity

We have previously described a method to measure the capacity of mutated ANP32 proteins to act as pro-viral factors for influenza polymerase activity. This is achieved by exogenous expression of the cloned mutants in human eHAP cells lacking ANP32A and ANP32B (dKO), in which influenza polymerase is unable to function in absence of complementing ANP32 (Staller et al. 2019; Peacock et al. 2020). Here we expressed each natural ANP32A or ANP32B variant and tested its ability to support reconstituted polymerases from a 2009 pandemic H1N1 isolate (A/England/195/2009; hereafter ‘Eng/195’) or a seasonal H3N2 virus (A/Victoria/3/75; hereafter ‘Vic/75’) in minigenome reporter assays (Figure 2).

**Figure 2.**
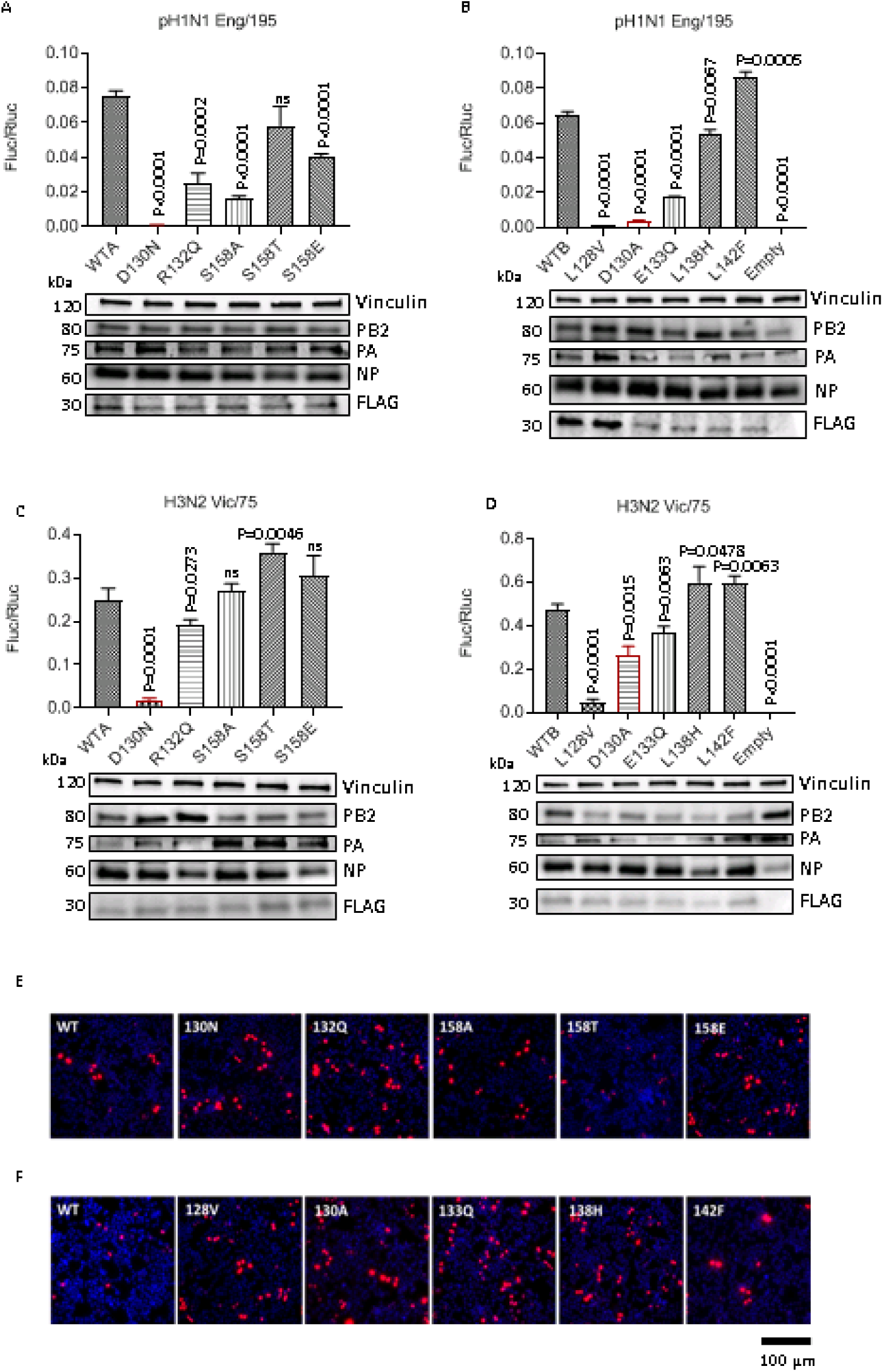
ANP32A and B mutant proteins show variable capacity to rescue IAV FluPol activity. (A-D) Minigenome reporter assays in eHAP dKO cells with co-transfected FLAG-tagged ANP32A (A and C) or ANP32B variants (B and D) with either pH1N1 Eng/195 (A and B) or H3N2 Vic/75 (C and D) RNP components PB1, PB2, PA and NP, pPolI-firefly luciferase minigenome reporter, and *Renilla* luciferase control in a 2:2:1:4:2:2 ratio. Data show mean (SD) of firefly activity normalized to *Renilla* and analysed by t test from one representative repeat (n = 3 independent triplicate experiments). Accompanying Western blots show expression of the FLAG-tagged ANP32 constructs alongside selected RNP components and endogenous vinculin loading control. (E-F) Immunofluorescence analysis showing nuclear localisation of FLAG-tagged ANP32A (E) or ANP32B (F) constructs, detected with anti-FLAG primary antibody and Alexa Fluor-568 anti-mouse conjugate and counterstained with DAPI.

Compared with wildtype ANP32A (WTA), ANP32A-D130N did not support Eng/195 polymerase activity at all, while R132Q and S158A substitutions had significantly reduced capacity to support FluPol activity (Figure 2A). An artificial mutant with the phosphomimic S158E, also had reduced capacity to support Eng/195 polymerase. In contrast, ANP32A-S158T supported FluPol to an extent similar to wildtype ANP32A. Compared with wildtype ANP32B (WTB), ANP32B-D130A had a large deleterious effect on the support for Eng/195 polymerase activity, as did the leucine to valine substitution at position 128 (ANP32B-L128V) (Figure 2B). ANP32B-E133Q was also significantly less able to support Eng/195 FluPol activity. Substitutions of the leucines at positions 138 and 142 to histidine (L138H) and phenylalanine (L142F), respectively, did not compromise the ability of the mutant ANP32B proteins to support Eng/195 polymerase activity.

In general the effects of natural variation in ANP32 proteins was similar for Vic/75 polymerase. ANP32A-D130N did not support Vic/75 polymerase activity, compared with WTA (Figure 2C), but R132Q and S158A substitutions had a smaller effect on Vic/75 FluPol activity than on Eng/195 FluPol activity. The S158T substitution and the phosphomimic S158E were as capable of supporting Vic/75 FluPol activity as WTA. As seen with Eng/195 polymerase, the ANP32B-L128V substitution was unable to rescue Vic/75 polymerase activity, but the D130A mutation had less detrimental effect and resulted in only a <2-fold reduction in Vic/75 polymerase activity (Figure 2D). This is in contrast to Eng/195 polymerase activity complemented with ANP32B-D130A, where a reduction of >17-fold was observed (compare Figures 2B and D). All in all Eng/195 polymerase, which has circulated in humans since 2009, was more sensitive to alterations in ANP32 than Vic/75 polymerase, which has replicated in human cells for over 40 years. The observed differences in FluPol activity were not explained by differences in expression of the FLAG-tagged ANP32 constructs (Western blots accompanying Figures 2A-D), nor by impaired nuclear localisation of the mutant proteins (Figures 2E and F).

### ANP32 position 130 mutants show impaired binding to FluPol

Using a split luciferase complementation assay previously developed in our laboratory (Mistry et al., 2020) (Figure 3A) we next assessed whether mutations at amino acid 130 affected interaction between ANP32 and FluPol. We and others have previously shown that murine ANP32A, which naturally harbours 130A, does not support influenza virus polymerase in minireplicon assays. Substituting 130A in human ANP32A greatly reduced FluPol activity, while introducing 130D in mouse ANP32A rescued its capacity to support FluPol activity (Staller et al., 2019; H. Zhang et al., 2019). Indeed, ANP32B KO mice, but not ANP32A KO mice, showed reduced viral loads and mortality when infected with H3N2 or H5N1 influenza A virus (Beck et al., 2020). Poor replication of H7N9 viruses in ANP32A knockout mice has also been described however (Liang et al., 2019). Interestingly, interaction of murine ANP32A with FluPol was reduced 2.5-fold compared with mouse ANP32B, which does have pro-influenza virus activity, and 3.2-fold compared with wildtype human ANP32A (Figure 3B).

**Figure 3.**
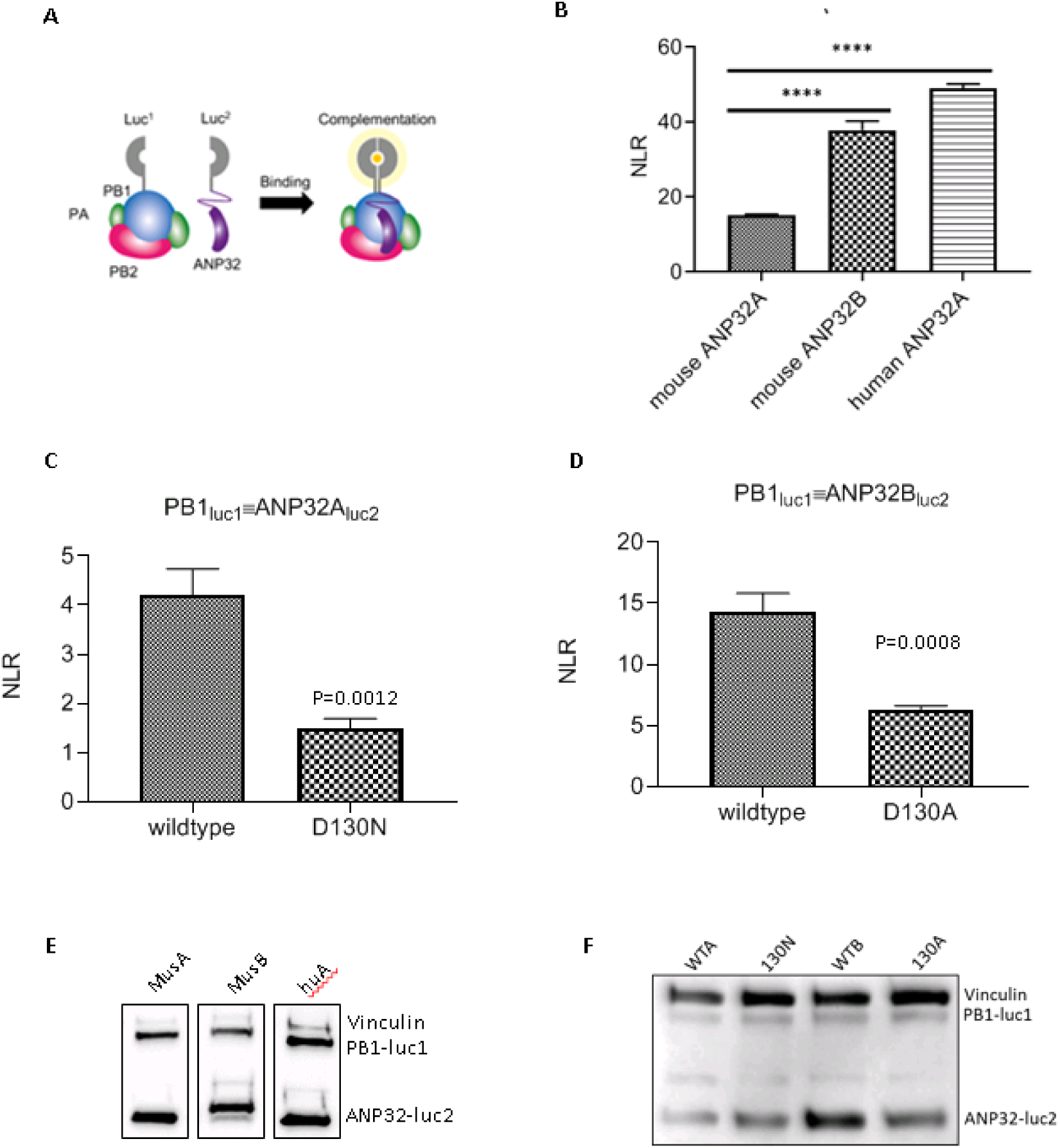
ANP32 position 130 mutants show impaired binding to FluPol. (A) Schematic of split luciferase complementation assay (adapted from Mistry et al. 2019). (B) Interaction between mouse ANP32A, mouse ANP32B or human ANP32A and Vic/75 FluPol. Accompanying Western blot analysis shows expression of luc1-tagged PB1 and luc2-tagged ANP32 constructs and vinculin loading control. (C-D) Interaction between naturally occurring human ANP32 variants ANP32A-D130N (C) and ANP32B-D130A (D) and Vic/75 FluPol. Split luciferase complementation assays were carried by transfecting equal amounts (15 ng) of pCAGGS expression plasmids encoding Vic/75 polymerase components PB1-luc1, PB2 and PA, as well as the indicated ANP32-luc2 construct into 293T cells. 24 hours post-transfection cells were lysed and luminescence was measured. Total amount of plasmid (in ng) was kept constant by using empty pCAGGS. Normalised luminescence ratio (NLR) was obtained as described in methods by dividing luminescence in the experimental condition (tagged PB1 + tagged ANP32) by the sum of the luminescence measured in the control conditions (i.e. background interaction of unbound luc1 with ANP32-luc2, and unbound luc2 with PB1-luc1, respectively). Data shown are mean (SD) representative of 3 independent triplicate experiments; statistical analysis in (B) by one-way ANOVA and in (C) and (D) by t-test. ****, P < 0.0001. (E) Western blot showing expression of luc1-tagged H3N2 Vic/75 PB1 and luc2-tagged ANP32 constructs alongside vinculin loading control.

The naturally occurring human ANP32 variants ANP32A-D130N and ANP32B-D130A also showed reduced binding to FluPol (Figures 3C and D). The luciferase signal indicating binding of ANP32A-D130N to FluPol was reduced almost 3-fold, relative to wildtype ANP32A and the signal indicating FluPol interaction of ANP32B-D130A was reduced >2-fold compared with wild type ANP32B. These differences in binding affinity were not due to differential expression of the PB1-luc1 and ANP32-luc2 constructs (Figures 3E and F).

### ANP32 position 130 mutants exert dominant-negative effects

To investigate the functional consequence of the ANP32B-D130A variant, we recapitulated heterozygous or homozygous ANP32B-D130A variant genotypes by exogenous expression of the mutated proteins in CRISPR-edited eHAP cells that lack ANP32B expression (BKO) (Staller et al., 2019) (Figure 4A). Minigenome assays were performed with reconstituted pH1N1 Eng/195 polymerase complemented by transient transfection with increasing amounts of either wildtype ANP32B (homozygous wildtype; blue bars), ANP32B-D130A (homozygous mutant; red bars) or a 1:1 ratio of both (heterozygous mutant; purple bars). We found that adding increasing amounts of wildtype ANP32B, essentially mimicking the homozygous wildtype genotype (blue bars), had no effect on FluPol activity compared with FluPol supported by ANP32A alone (grey bar). This is presumably due to the presence of wildtype ANP32A in the BKO cells, since ANP32A and ANP32B serve redundant roles in supporting FluPol. In contrast, polymerase activity decreased significantly when ANP32B-D130A rather than wildtype ANP32B was co-transfected, and this effect was more dramatic the more variant that was expressed (red bars). This suggests paralogue interference of ANP32B-D130A over wildtype ANP32A, which would otherwise support polymerase activity effectively (grey bar). Co-transfecting increasing amounts of wildtype and mutant ANP32B in a 1:1 ratio, recapitulating the heterozygous mutant genotype (purple bars), also resulted in a significant drop in polymerase activity. This suggests that ANP32B-D130A also exerts a dominant-negative effect over wildtype ANP32B. These observations were unrelated to expression levels of the FLAG-tagged ANP32 constructs (accompanying Western blots). Statistical significance of the differences in Eng/195 polymerase activity between conditions is shown in the accompanying Table.

**Figure 4.**
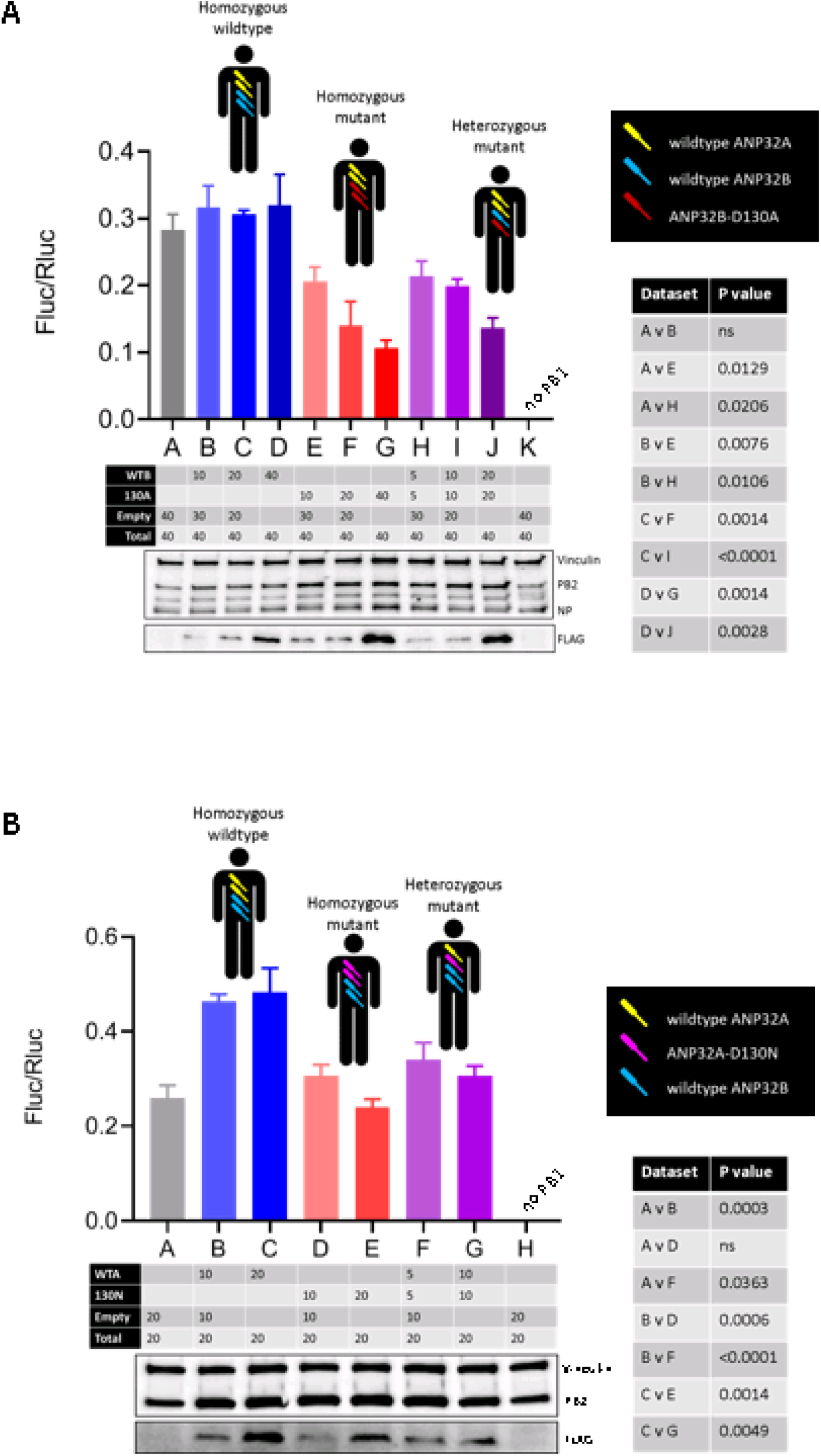
ANP32B-D130A exerts dominant-negative effects on pro-viral function of wildtype ANP32B and ANP32A. (A) Minigenome reporter assay in eHAP cells lacking ANP32B expression (BKO) with 2009 pH1N1 Eng/195 RNP components PB1, PB2, PA and NP, pPolI-firefly luciferase minigenome reporter, and *Renilla* luciferase control, in a 2:2:1:4:2:2 transfection ratio with 20 ng pCAGGS-PB1 and 10 ng pCAGGS-PA per well (∼1.0 × 10^5^ cells). Complementation by co-transfection of indicated combinations of wildtype ANP32B, ANP32B-D130A, or empty pCAGGS plasmid (in ng) is shown in the Table. Total amounts of transfected plasmid DNA were kept constant across conditions by supplementing with empty pCAGGS plasmid. Data shown are mean (SD) of firefly activity normalized to *Renilla* from one representative biological repeat (n = 3 independent triplicate experiments); statistical analysis in the accompanying table was carried out by t test. Accompanying Western blot shows expression of the FLAG-tagged ANP32 constructs alongside RNP components PB2 and NP, and vinculin loading control. Cartoon figures show the wildtype and mutant genotypes found in carriers of SNV rs182096718. All three genotypes have two alleles of wildtype *Anp32A* (yellow). Homozygous ANP32B wildtype individuals carry two wildtype *Anp32B* alleles (blue); homozygous mutant carriers have two alleles encoding ANP32B-D130A (red), and finally heterozygous mutant carriers have one wildtype and one variant *Anp32B* allele. (B) Minigenome reporter assay in eHAP cells lacking ANP32A expression (AKO) with 2009 pH1N1 Eng/195 RNP components PB1, PB2, PA and NP, pPolI-firefly luciferase minigenome reporter, and *Renilla* luciferase control, in a 2:2:1:4:2:2 transfection ratio. Co-transfected wildtype ANP32A, ANP32A-D130N and Empty pCAGGS plasmid are indicated in the Table, as in (A). Accompanying Western blot shows expression of the FLAG-tagged ANP32 constructs alongside RNP component PB2 and a vinculin loading control. Cartoon figures show the wildtype and mutant genotypes found in carriers of SNV rs771143708. All three genotypes have two alleles of wildtype *Anp32B* (red). Homozygous ANP32A wildtype individuals carry two wildtype *Anp32A* alleles (yellow); hypothetical homozygous mutant carriers have two alleles encoding ANP32A-D130N (pink), and finally heterozygous mutant carriers have one wildtype and one variant *Anp32A* allele.

We next investigated whether similar dominant-negative effects were exerted by the rare ANP32A-D130N variant (Figure 4B). In eHAP cells lacking ANP32A expression (AKO) pH1N1 Eng/195 polymerase activity increased significantly when wildtype ANP32A was provided by transient co-transfection (blue bars), compared with FluPol activity supported by ANP32B alone (grey bar). In contrast, decreased polymerase activity was evident when either ANP32A-D130N alone (red bars) or wildtype and mutant ANP32A in a 1:1 ratio (purple bars) were co-transfected. This suggests that, like ANP32B-D130A, ANP32A-D130N is dominant-negative over wildtype ANP32A, and exerts paralogue interference over wildtype ANP32B. Together these data show that different substitutions of the canonical aspartate at position 130 of both pro-influenza viral human ANP32 proteins can have similar phenotypic effects and impact influenza proviral activity even in the heterozygote genotypes.

### IAV polymerase activity and replication are attenuated in ANP32B-D130A mutant cells

To further probe the potential significance of the relatively common ANP32B-D130A variant, we generated a cell line with the wildtype *Anp32A* / mutant *Anp32B* (D130A) genotype by CRISPR/Cas9 genome editing, using human codon-optimised SpCas9 and a single-stranded DNA (ssODN) homology-directed repair template. We also generated a cell line lacking the entire pro-viral 128-130 loop (Δ128-130) (Figure 5A). We selected an unsuccessfully edited clone to serve as a negative control, and carried out Western blotting analysis to ensure endogenous wildtype ANP32A and wildtype or mutant ANP32B were expressed in control and mutant cell lines (Figure 5B). Using minigenome reporter assays, we found that reconstituted IAV polymerases from Eng/195 (Figure 5C) or Vic/75 (Figure 5D) were significantly less active in edited ANP32B-D130A and Δ128-130 cells, compared with control cells. As previously described (Staller et al. 2019), polymerase activity was not attenuated in BKO cells, again suggesting paralogue interference of mutant ANP32B over wildtype ANP32A. In other words, IAV FluPol activity was higher in cells completely lacking ANP32B expression than in cells expressing the D130A mutant. As described previously (Staller et al., 2019), no polymerase activity was seen in cells lacking both ANP32A and ANP32B (dKO), nor in the negative control conditions where the polymerase subunit PB2 was lacking (2P). Western blotting analysis showed the observed effects were not due to differential expression of transfected RNP components.

**Figure 5.**
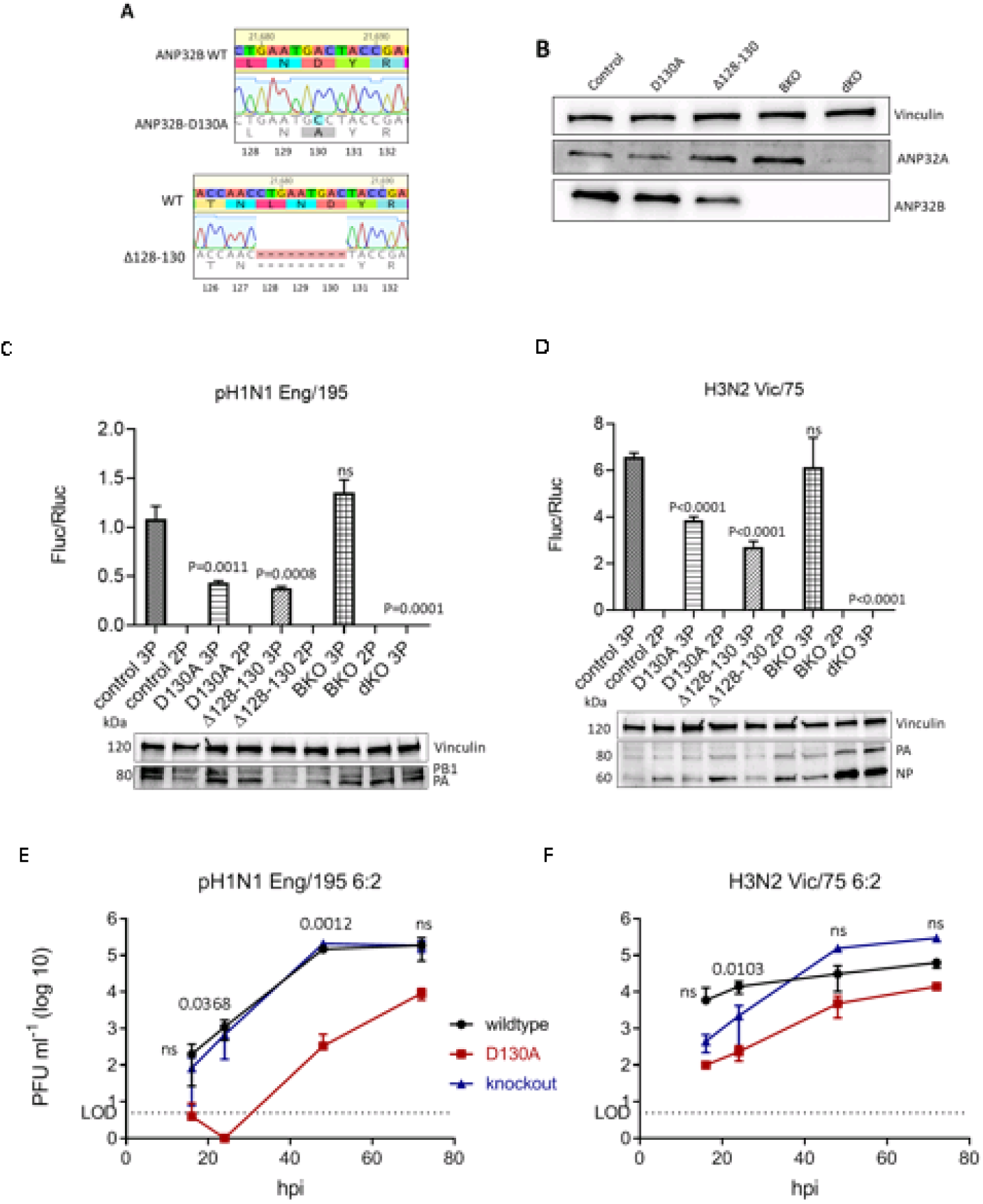
FluPol activity and virus replication are attenuated in ANP32B-D130A mutant cells. (A) Genotypic profiles of CRISPR-edited monoclonal eHAP cell lines expressing ANP32B-D130A (top) or ANP32B lacking the pro-influenza virus 128-130 loop (Δ128-130). (B) Western blotting analysis showing expression of endogenous ANP32A and ANP32B, alongside vinculin loading control, in CRISPR-edited monoclonal control, ANP32B-D130A and ANP32B-Δ128-130 cell lines, as well as the previously established ANP32B knockout (BKO) and ANP32A and B double knockout (dKO) cell lines. (C-D) Minigenome reporter assays in CRISPR-edited cell lines with pH1N1 Eng/195 (C) or H3N2 Vic/75 (D) RNP components PB1, PB2, PA and NP, pPolI-firefly luciferase minigenome reporter, and *Renilla* luciferase control, in a 2:2:1:4:2:2 ratio. Data show mean (SD) of firefly activity normalized to *Renilla* and analysed by t-test from one representative repeat (n = 3 independent triplicate experiments). Accompanying Western blots show expression of RNP components PB1, PA and NP alongside vinculin loading control. 3P, FluPol subunits PB1, PB2 and PA; 2P, negative control lacking FluPol subunit PB2. (E-F) CRISPR-edited control (wildtype; black), mutant ANP32B-D130A (D130A; red), and ANP32B knockout (blue) monoclonal cell lines were infected with either England/195 or Victoria/75 6:2 reassortant viruses with PR8 hemagglutinin (HA) and neuraminidase (NA) external genes at MOI = 0.005, and incubated at 37°C in the presence of 1 µg/ml trypsin to allow multicycle replication. Supernatants were harvested at the indicated times post-infection and pfu/ml established by plaque assay on MDCK cells. LOD (dotted line) denotes the limit of detection in the plaque assays. Shown are representative data from one of two independent infection experiments carried out in triplicate. P values were calculated per time point by t test and represent differences between control and mutant cell lines.

Next we infected mutant (ANP32B-D130A), ANP32B knockout, or control cells at low-MOI (0.005) with 6:2 recombinant influenza A viruses containing internal genes from pH1N1 Eng/195 and H3N2 Vic/75, with the neuraminidase (NA) and haemagglutinin (HA) external genes of the laboratory-adapted H1N1 PR/8 virus. Eng/195 6:2 virus replication was severely attenuated in mutant D130A cells, in comparison with either control or BKO cells (Figure 5E). Plaques were below the level of detection (LOD) 16 and 24 hours post-infection and still 3 logs lower than control cells 48 hours after infection (P=0.0012). Vic/75 6:2 infectious virus production in mutant cells was also lower than in control cells (Figure 5F) but less affected than Eng/195. Both viruses replicated to higher titres in cells that completely lacked ANP32B (BKO) than in cells expressing the ANP32B-D130A mutant, suggesting paralogue interference of ANP32B-D130A over wildtype ANP32A also in an infectious virus context.

## DISCUSSION

Here, we used publicly available databases to perform a biased screen for naturally occurring single nucleotide variants in human *Anp32A* and *Anp32B*, under the hypothesis that some mutations may affect susceptibility to influenza virus infection. One of the mutations, which translates to an aberrant ANP32B-D130A protein, was highly enriched in carriers of Latino descent. Although most carriers of this SNV (>1,200) are heterozygous for the mutation, 25 homozygotes have been reported so far in the gnomAD database. Almost all carriers occur in the Hispanic/Latino subpopulation. We generated this naturally occurring homozygous genotype by CRISPR/Cas9 genome editing in low-ploidy human eHAP cells, and found that IAV FluPol activity and virus replication were significantly attenuated in the mutant cell line, compared to wildtype control cells or cells lacking ANP32B. This effect was most pronounced with the 2009 pandemic H1N1 strain Eng/195. Importantly, we demonstrate that the mutant ANP32B-D130A protein exerts a dominant-negative effect over wildtype ANP32B, as well as interference over the functionally redundant paralogue ANP32A. This suggests that despite the redundancy in pro-influenza viral function of ANP32A and B, even in heterozygotes there may be some consequence of this genetic variation to outcome of virus infection. We provide the first example of a single nucleotide variant in the coding region of a human gene that might confer some protection against influenza virus infection.

Although we and others have previously reported redundancy in reconstituted systems for ANP32A and B in their support of FluPol activity (Staller et al., 2019; H. Zhang et al., 2019), the data presented here imply that deleterious mutations in one ANP32 protein might exert dominant-negative effects on the pro-viral activity of the other, a phenomenon known as paralogue interference. Poor interaction with FluPol of one paralogue might be expected to make no difference when both paralogues are present in the cell, but clearly this does not seem to be the case.

Recent structural data suggest that ANP32 proteins assist influenza polymerase by stabilizing a dimer between the replicating FluPol trimer, FluPol^R^, and a second trimer that might be recruited as an encapsidating complex, FluPol^E^ (Carrique et al. 2020, *in press*). To explain the dominant-negative effects we observe here, as well as the compromised replicative activity, we propose a model (Figure 6) in which the mutant ANP32B-D130A binds competitively to FluPol^R^, via the unaltered N-terminal portion of the LRR domain, but is unable to recruit FluPol^E^ due to the lack of acidity at position 130. Consequently, nascent viral RNA generated by FluPol^R^ is not transferred effectively to FluPol^E^, and replication is compromised. Furthermore, recruitment of ANP32B-D130A to FluPol^R^ occupies the binding site for wildtype ANP32A or ANP32B proteins, explaining the paralogue interference and dominant-negative effects we observe, respectively.

**Figure 6.**
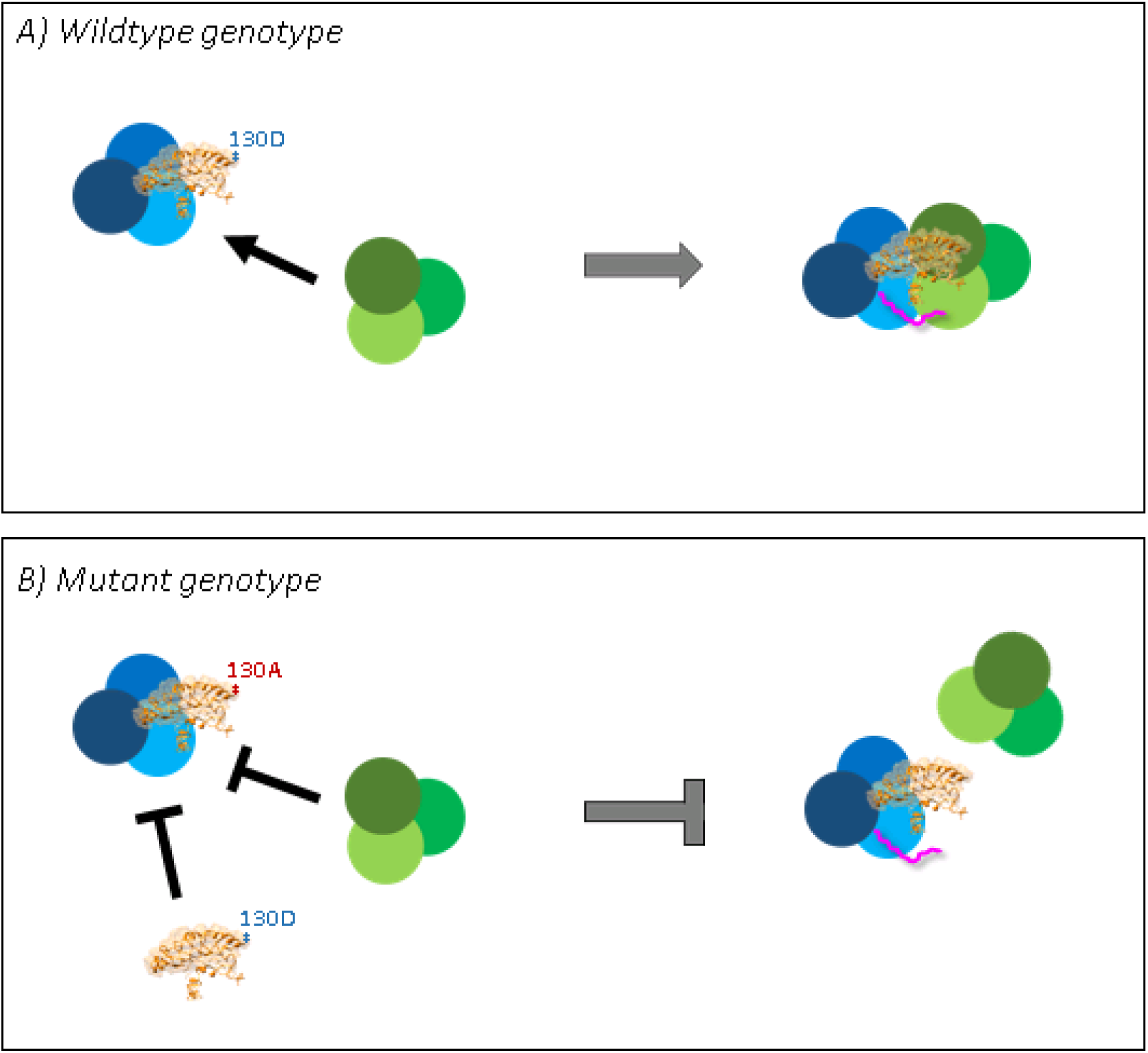
Mutation at ANP32 position 130 results in impaired formation of replication-competent FluPol dimer. Cartoon explaining why FluPol activity is impaired in cells harbouring ANP32B-D130A. Wildtype (130D) ANP32B (or ANP32A) acts by binding to a viral promoter-bound replicating FluPol (FluPol^R^, in blue) via the N-terminal region of the LRR domain, while the C-terminal LRR, harbouring the surface-exposed acidic residue 130D, is used to recruit the encapsidating FluPol (FluPol^E^, in green), via direct interaction with basic residues on the PB1 subunit of FluPol^E^ (A). In the context of a properly formed FluPol^R^-ANP32-FluPol^E^ complex the nascent RNA product replicated by FluPol^R^ (in pink) transfers to FluPol^E^ and replication proceeds efficiently. The mutant ANP32B-D130A can still bind efficiently – and therefore competitively – to FluPol^R^, as the N-terminal portion of the LRR is unaffected (B), but because there is now an alanine in place of the acidic aspartate at position 130, recruitment of FluPol^E^ is impaired. Furthermore, binding of ANP32B-D130A to FluPol^R^ prevents recruitment of wildtype ANP32A or ANP32B, explaining the paralogue interference and dominant-negative effects, respectively. Thus, in the presence of ANP32B-D130A, the replication-competent FluPol^R^-ANP32-FluPol^E^ complex is not formed as efficiently, and therefore replication is compromised.

It is important to bear in mind that the outcome of genetic variation will depend on gene dosing in relevant tissues. The genotype-tissue expression (GTEx) portal uses whole genome sequencing and RNA-Seq data to estimate gene expression in 54 non-diseased human tissue types from nearly 1,000 individuals (Lonsdale et al., 2013). ANP32A and ANP32B are both clearly detectable in healthy lung tissue, at a median transcripts per kilobase million (TPM) of 42 and 201, respectively. There are thus 4.8-fold more ANP32B transcripts than ANP32A transcripts in the lung, suggesting that mutations that impair pro-viral function of ANP32B, such as ANP32B-D130A, might have significant effects on virus replication in the target tissue. Moreover, ANP32B is more stable than ANP32A, judging from half-life measurements of both paralogues (Fries et al., 2007).

A caveat of the work presented here is the use of human eHAP cells where (primary) respiratory epithelium would have been preferable. We selected eHAP cells for their low ploidy, which renders CRISPR/Cas9 genome editing more straightforward, in the knowledge that more commonly used laboratory cell lines like A549 or HEK-293 derivatives are polyploid (Giard et al., 1973; Graham, Smiley, Russell, & Nairn, 1977; Lin et al., 2014). We and others have, however, shown previously that eHAP cells are a suitable model for influenza virus infection (Jason S. Long et al., 2019; Mistry et al., 2020; Thomas. P. Peacock et al., 2020; Thomas P. Peacock et al., 2020; Staller et al., 2019) (Carrique et al, 2020, *in press*). A drawback of using a low-ploidy cell line is that heterozygous genotypes cannot easily be generated. Thus far we have been unable to trace carriers of the variants but future work using diploid respiratory epithelial cells that reflect the heterozygous genotype could indicate whether true heterozygotes, like the reconstituted cells we used here, also display reduced influenza susceptibility.

In conclusion, we have provided the first example of a single nucleotide variant in the coding region of a human gene that may offer carriers some protection against influenza virus. This work has the potential to inform future intervention.

## MATERIALS AND METHODS

### Cell culture

Human eHAP cells (Horizon Discovery) were cultured in Iscove’s modified Dulbecco’s medium (IMDM; Thermo Fisher) supplemented with 10% fetal bovine serum (FBS; Labtech), 1% nonessential amino acids (NEAA; Gibco), and 1% penicillin/streptomycin (Invitrogen). Human embryonic kidney (293T) cells (ATCC) and Madin-Darby canine kidney (MDCK) cells (ATCC) were maintained in Dulbecco’s modified Eagle’s medium (DMEM; Invitrogen) supplemented with 10% FBS, 1% NEAAs, and 1% penicillin/streptomycin. All cells were maintained at 37°C in a 5% CO^2^ atmosphere.

### Plasmids and cloning

Human FLAG-tagged pCAGGS-ANP32A and ANP32B expression plasmids have been described (Staller et al., 2019). Mutant proteins were cloned from these plasmids by overlapping touchdown PCR, using primers CCAACCTGAATAACTACCGCGAGAAC and GTTCTCGCGGTAGTTATTCAGGTTGG (ANP32A-D130N), GAATGACTACCAAGAGAACGTGTTC and GAACACGTTCTCTTGGTAGTCATTC (ANP32A-R132Q), GAGGCCCCTGATGCTGACGCCGAGG and CCTCGGCGTCAGCATCAGGGGCCTC (ANP32A-S158A), GAGGCCCCTGATACTGACGCCGAGGGC and GCCCTCGGCGTCAGTATCAGGGGCCTC (ANP32A-S158T), GTGACAAACGTGAATGACTATCGG and CCGATAGTCATTCACGTTTGTCAC (ANP32B-L128V), CAAACCTGAATGCCTATCGGGAGAGC and GCTCTCCCGATAGGCATTCAGGTTTG (ANP32B-D130A), GACTATCGGCAGAGCGTGTTTAAG and CTTAAACACGCTCTGCCGATAGTC (ANP32B-E133Q), GAGAGCGTGTTTAAGCACCTGCCACAGCTG and CAGCTGTGGCAGGTGCTTAAACACGCTCTC (ANP32B-L138H), GTTGCTGCCACAGTTTACTTATCTCGA and TCGAGATAAGTAAACTGTGGCAGCAAC (ANP32B-L142F). Expression plasmids encoding H3N2 Vic/75 and pH1N1 RNP components PB1, PB2, PA and NP have been described (Staller et al., 2019), as have pCAGGS-ANP32A_luc2C_, pCAGGS-luc1, pCAGGS-luc2, and H5N1 50-92 PB1_luc1C_ (Jason S. Long et al., 2019; Mistry et al., 2020). pPolI reporter plasmid containing firefly luciferase flanked by IAV-specific promoters, and pCAGGS-*Renilla* luciferase transfection / cellular transcription control have been previously described (Staller et al., 2019). Expression plasmids pCAGGS-ANP32B_luc2C_, ANP32A-D130N_luc2C_, ANP32B-L128V_luc2C_, and ANP32B-D130A_luc2C_ were cloned by overlapping touchdown PCR. All plasmid constructs were verified by Sanger sequencing and analyzed in Geneious prime 2019.

### CRISPR/Cas9 genome editing

Guide RNA GCACTCTCTCGGTAGTCATTC was designed manually against the protospacer sequence in exon 4 of *Anp32B* to target DNA endonuclease SpCas9, expressed from Addgene plasmid # 62988 (PX459), to the target nucleotide (Supplementary Figure 3A). The guide RNA itself was cloned into Addgene plasmid # 80457 (pmCherry_gRNA). A custom-designed 88 base ssODN (single strand DNA) homology-directed repair template, harbouring the point mutation (cytosine in bold) and a silent PAM mutation (thymidine in italic script) – GAAAAGCCTGGACCTCTTTAACTGTGAGGTTACCAA*T*CTGAATG**C**CTACCGAGAGAGTG TCTTCAAGCTCCTGCCCCAGCTTACCTAC – was obtained from Integrated DNA Technologies (IDT). Equal amounts of PX459 and pmCherry_gRNA (total 1 µg), and 1.0 µl 10 µM ssODN template, were transfected by electroporation into approximately 400,000 eHAP cells using the Neon™ transfection system (Invitrogen). Cells and DNA were mixed into a 10 µl volume of suspension buffer, which was subjected to a single 1,200 Volt pulse for a duration of 40 ms. Cells were incubated at 37°C for 24 hours in IMDM growth medium without antibiotics. A fluorescence-activated cell sorter (FACS) Aria IIIU (BD Biosciences) with an 85-µm nozzle was used to sort cells expressing mCherry (550-650 nm emission) into 96-well plates containing growth medium. Single cells were grown out into monoclonal populations over a period of 10 to 14 days. Total genomic DNA was extracted using the Purelink™ Genomic DNA Mini Kit (Invitrogen) and amplified by touchdown PCR to generate a 1,561-base pair fragment of the edited locus (primers TACCTCTGCCCTCTCAATCTCT and ACGCACACAAACACACACTATT). PCR products were then incubated at 65°C for 30 minutes in the presence of BsmI restriction enzyme (NEB). The resulting DNA fragments were separated by 1.5% agarose gel electrophoresis. Potentially successfully edited clones were verified by Sanger sequencing (primers TAAAGACCGCTTGATACCCAGG and TGAGGCTGAGTGGGTAGTGG) and analysed in Geneious prime 2019.

### Minigenome reporter assays

In order to measure influenza virus polymerase activity, pCAGGS expression plasmids encoding H3N2 Vic/75 or pH1N1 Eng/195 PB1 (0.02 µg), PB2 (0.02 µg), PA (0.01 µg), and NP (0.04 µg) were transfected into ∼100,000 eHAP cells using Lipofectamine 3000 (Thermo Fisher) at ratios of 2 µl P3000 reagent and 3 µl Lipofectamine 3000 reagent per µg plasmid DNA. As a reporter construct, we transfected 0.02 µg pPolI-luc, which encodes a minigenome containing a firefly luciferase reporter flanked by influenza A virus promoter sequences. pCAGGS-*Renilla* luciferase (0.02 µg) was co-transfected as a transfection and toxicity control. Amounts of co-transfected ANP32-FLAG constructs were 0.04 ug (equal to pCAGGS-NP) unless otherwise specified in the Figure legends. Twenty-four hours after transfection, cells were lysed in 50 µl passive lysis buffer (Promega) for 30 minutes at room temperature with gentle shaking. Bioluminescence generated by firefly and *Renilla* luciferases was measured using the dual-luciferase system (Promega) on a FLUOstar Omega plate reader (BMG Labtech).

### Split luciferase complementation assay

pCAGGS expression plasmids encoding H3N2 Vic/75 PB1-luc1, PB2, PA, and the indicated ANP32-luc2 construct were transfected into ∼100,000 293T cells at a ratio of 1:1:1:1 (15 ng per well). Control conditions contained pCAGGS-luc1 and untagged PB1, or pCAGGS-luc2 and untagged ANP32A, respectively, with all other components remaining constant. Empty pCAGGS plasmid was used to ensure total transfected DNA was equal across conditions. Twenty-four hours after transfection, cells were lysed in 50 µl *Renilla* lysis buffer (Promega) for 1 h at room temperature with gentle shaking (*Gaussia* and *Renilla* luciferase share the same substrate). Bioluminescence generated by *Gaussia* luciferase was measured using the *Renilla* luciferase kit (Promega) on a FLUOstar Omega plate reader (BMG Labtech). Normalized luminescence ratios (NLR) were calculated by dividing the signal from the potential interacting partners by the sum of the two controls, as described (Mistry et al., 2020).

### Western blotting

At least 250,000 cells were lysed in buffer containing 50 mM Tris-HCl (pH 7.8; Sigma-Aldrich), 100 mM NaCl, 50 mM KCl, and 0.5% Triton X-100 (Sigma-Aldrich), supplemented with a cOmplete EDTA-free protease inhibitor cocktail tablet (Roche) and prepared in Laemmli 4x buffer (Bio-Rad) after protein concentration had been established by spectrophotometry (DeNovix DS-11 FX+ spectrophotometer). Equal amounts of total protein (20-60 µg per lane) was resolved by SDS-PAGE using Mini Protean TGX precast gels 4% to 20% (Bio-Rad). Immunoblotting by semi-dry transfer (Bio-Rad Trans-Blot SD semidry transfer cell) onto nitrocellulose membranes (Amersham Protran Premium 0.2 µm NC; GE Healthcare) was carried out using the following primary antibodies: rabbit α-vinculin (catalogue number ab129002, 1/2,000; Abcam), rabbit α-ANP32A (catalogue number ab51013, 1/500; Abcam), rabbit α-ANP32B (10843-1-AP, 1/1,000; Proteintech), mouse α-FLAG (catalogue number F1804, 1/500; Sigma-Aldrich), mouse α-NP (catalogue number ab128193, 1/1,000; Abcam), rabbit α-PB1 (catalogue number GTX125923, 1/500; GeneTex), rabbit α-PA (catalogue number GTX118991, 1/500; GeneTex), and rabbit α-IAV PB2 (catalogue number GTX125926, 1/2,000; GeneTex). The following secondary antibodies were used: sheep α-rabbit horseradish peroxidase (HRP) (catalogue number AP510P, 1/10,000; Merck) and goat α-mouse HRP (STAR117P, 1/5,000; AbD Serotec). Protein bands were visualized by chemiluminescence using SuperSignal™ West Femto substrate (Thermofisher Scientific) on a Fusion-FX imaging system (Vilber Lourmat).

### Immunofluorescence microscopy

Approximately 100,000 eHAP cells were cultured on sterilised glass coverslips and transfected as per minigenome reporter assay protocol. Twenty-four hours after transfection, cells were fixed in 4% paraformaldehyde and permeabilized in 0.2% Triton X-100. FLAG-tagged ANP32 constructs were visualised with primary antibody mouse α-FLAG (F1804; 1/200; Sigma) for 2 hours at 37°C in a humidified chamber. Cells were incubated with secondary antibody goat α-mouse Alexa Fluor-568 (1/200; Life Technologies) for 1 hour at 37°C in a humidified chamber, and counterstained with DAPI. Coverslips were mounted on glass slides using Vectashield mounting medium (H-1000-10; Vector Laboratories). Cells were imaged with a Zeiss Cell Observer widefield microscope with ZEN Blue software, using a Plan-Apochromat x100 1.40-numerical aperture oil objective (Zeiss), an Orca-Flash 4.0 complementary metal-oxide semiconductor (CMOS) camera (frame, 2,048 × 2,048 pixels; Hamamatsu), giving a pixel size of 65 nm, and a Colibri 7 light source (Zeiss). Channels acquired and filters for excitation and emission were 4’,6-diamidino-2-phenylindole (DAPI) (excitation [ex], 365/12 nm, emission [em] 447/60 nm), and TexasRed (ex 562/40 nm, em 624/40 nm). All images were analyzed and prepared with Fiji software.

### Influenza virus infection

500,000 CRISPR/Cas9-modified monoclonal eHAP cells were infected with ∼2,500 plaque-forming units (PFU) H3N2 Vic/75 6:2 or pH1N1 Eng/195 6:2 virus diluted in 200 µl serum-free IMDM for 1 hour at 37°C (MOI 0.005) to allow virus to adsorb and enter the cells. The inoculum was removed and cells were incubated in room-temperature phosphate-buffered saline / HCl at pH 3.0 for 3 minutes to inactivate residual virus. Cells were incubated at 37°C in serum-free cell culture medium (IMDM) supplemented with 1 µg/ml L-1-tosylamide-2-phenylethyl chloromethyl ketone (TPCK) trypsin (Worthington-Biochemical). Cell supernatants were harvested at indicated time points post-infection. Infectious titres were determined by plaque assay on MDCK cells. Virus infection assays were performed in triplicate on two separate occasions.

### Safety/biosecurity

All work with infectious agents was conducted in biosafety level 2 facilities, approved by the Health and Safety Executive of the United Kingdom and in accordance with local rules, at Imperial College London, United Kingdom.

### Structural modelling

Structural models of ANP32A and B were created using iTASSER structural prediction software (based primarily on huANP32B [GenBank accession number 2RR6A] and huANP32A [accession number 2JQDA], and 2JEOA). The three-dimensional structural models were visualized and created in UCSF Chimera. Amino acid residues affected by selected SNVs are highlighted in purple (ANP32A) or blue (ANP32B) stick format.

### Bioinformatics

Human genomic information was obtained using the following publicly available databases: gnomAD (https://gnomad.broadinstitute.org/); NCBI dbSNP (https://www.ncbi.nlm.nih.gov/snp/); ALSPAC (http://www.bristol.ac.uk/alspac/); TOPMed (https://www.nhlbiwgs.org/); 1000G (https://www.internationalgenome.org/); GO-ESP (https://esp.gs.washington.edu/drupal/)

## ACKNOWLEDGEMENTS

The authors wish to thank David Gaboriau for help with microscopy, and the St. Mary’s NHLI FACS core facility for support, instrumentation, and help with single cell sorting. The Facility for Imaging by Light Microscopy (FILM) at Imperial College London is partially supported by funding from the Wellcome Trust (grant 104931/Z/14/Z) and BBSRC (grant BB/L015129/1) E.S. was supported by an Imperial College President’s Scholarship. L.B. was supported by Wellcome Trust grant 209213/Z/17/Z. R.F. was supported by Wellcome Trust grant 200187/Z/15/Z. T.P.P. was supported by BBSRC grant BB/R013071/1. V.S.S. was supported by UKRI Future Leaders Fellowship MR/S032304/1. C.M.S. and W.S.B. were funded by Wellcome Trust grant 205100 and BBSRC grant BB/K002456/1.

## REFERENCES

Allen, E. K., Randolph, A. G., Bhangale, T., Dogra, P., Ohlson, M., Oshansky, C. M., … Thomas, P. G. (2017). SNP-mediated disruption of CTCF binding at the IFITM3 promoter is associated with risk of severe influenza in humans. Nat Med, 23(8), 975–983. doi: 10.1038/nm.4370

Beck, S., Zickler, M., Pinho dos Reis, V., Günther, T., Grundhoff, A., Reilly, P. T., … Gabriel, G. (2020). ANP32B Deficiency Protects Mice From Lethal Influenza A Virus Challenge by Dampening the Host Immune Response. Frontiers in Immunology, 11(450). doi: 10.3389/fimmu.2020.00450

Bravo Garcia-Morato, M., Calvo Apalategi, A., Bravo-Gallego, L. Y., Blazquez Moreno, A., Simon-Fuentes, M., Garmendia, J. V., … Rodriguez Pena, R. (2019). Impaired control of multiple viral infections in a family with complete IRF9 deficiency. J Allergy Clin Immunol, 144(1), 309-312.e310. doi: 10.1016/j.jaci.2019.02.019

Camacho-Zarco, A. R., Kalayil, S., Maurin, D., Salvi, N., Delaforge, E., Milles, S., … Blackledge, M. (2020). Molecular basis of host-adaptation interactions between influenza virus polymerase PB2 subunit and ANP32A. Nat Commun, 11(1), 3656. doi: 10.1038/s41467-020-17407-x

Carette, J. E., Raaben, M., Wong, A. C., Herbert, A. S., Obernosterer, G., Mulherkar, N., … Ruthel, G. (2011). Ebola virus entry requires the cholesterol transporter Niemann–Pick C1. Nature, 477(7364), 340–343.

Carrington, M., Dean, M., Martin, M. P., & O’Brien, S. J. (1999). Genetics of HIV-1 Infection: Chemokine Receptor Ccr5 Polymorphism and Its Consequences. Human Molecular Genetics, 8(10), 1939–1945. doi: 10.1093/hmg/8.10.1939

Chang, S., Sun, D., Liang, H., Wang, J., Li, J., Guo, L., … Liu, Y. (2015). Cryo-EM structure of influenza virus RNA polymerase complex at 4.3 Å resolution. Mol Cell, 57(5), 925–935. doi: 10.1016/j.molcel.2014.12.031

Chatzopoulou, F., Gioula, G., Kioumis, I., Chatzidimitriou, D., & Exindari, M. (2019). Identification of complement-related host genetic risk factors associated with influenza A(H1N1)pdm09 outcome: challenges ahead. Med Microbiol Immunol, 208(5), 631–640. doi: 10.1007/s00430-018-0567-9

Chen, K. Y., Santos Afonso, E. D., Enouf, V., Isel, C., & Naffakh, N. (2019). Influenza virus polymerase subunits co-evolve to ensure proper levels of dimerization of the heterotrimer. PLoS Pathog, 15(10), e1008034. doi: 10.1371/journal.ppat.1008034

Chen, Y., Zhou, J., Cheng, Z., Yang, S., Chu, H., Fan, Y., … Li, L. (2015). Functional variants regulating LGALS1 (Galectin 1) expression affect human susceptibility to influenza A(H7N9). Sci Rep, 5, 8517. doi: 10.1038/srep08517

Cheng, Z., Zhou, J., To, K. K., Chu, H., Li, C., Wang, D., … Yuen, K. Y. (2015). Identification of TMPRSS2 as a Susceptibility Gene for Severe 2009 Pandemic A(H1N1) Influenza and A(H7N9) Influenza. J Infect Dis, 212(8), 1214–1221. doi: 10.1093/infdis/jiv246

Ciancanelli, M. J., Abel, L., Zhang, S. Y., & Casanova, J. L. (2016). Host genetics of severe influenza: from mouse Mx1 to human IRF7. Curr Opin Immunol, 38, 109–120. doi: 10.1016/j.coi.2015.12.002

Ciancanelli, M. J., Huang, S. X., Luthra, P., Garner, H., Itan, Y., Volpi, S., … Casanova, J. L. (2015). Infectious disease. Life-threatening influenza and impaired interferon amplification in human IRF7 deficiency. Science, 348(6233), 448–453. doi: 10.1126/science.aaa1578

Esposito, S., Molteni, C. G., Giliani, S., Mazza, C., Scala, A., Tagliaferri, L., … Principi, N. (2012). Toll-like receptor 3 gene polymorphisms and severity of pandemic A/H1N1/2009 influenza in otherwise healthy children. Virol J, 9, 270. doi: 10.1186/1743-422x-9-270

Everitt, A. R., Clare, S., Pertel, T., John, S. P., Wash, R. S., Smith, S. E., … Wellcome Trust Sanger Institute, U. K. (2012). IFITM3 restricts the morbidity and mortality associated with influenza. Nature, 484(7395), 519–523. doi: 10.1038/nature10921

Fan, H., Walker, A. P., Carrique, L., Keown, J. R., Serna Martin, I., Karia, D., … Fodor, E. (2019). Structures of influenza A virus RNA polymerase offer insight into viral genome replication. Nature, 573(7773), 287–290. doi: 10.1038/s41586-019-1530-7

Fodor, E., & Te Velthuis, A. J. W. (2019). Structure and Function of the Influenza Virus Transcription and Replication Machinery. Cold Spring Harb Perspect Med. doi: 10.1101/cshperspect.a038398

Fries, B., Heukeshoven, J., Hauber, I., Gruttner, C., Stocking, C., Kehlenbach, R. H., … Chemnitz, J. (2007). Analysis of nucleocytoplasmic trafficking of the HuR ligand APRIL and its influence on CD83 expression. J Biol Chem, 282(7), 4504–4515. doi: 10.1074/jbc.M608849200

Giard, D. J., Aaronson, S. A., Todaro, G. J., Arnstein, P., Kersey, J. H., Dosik, H., & Parks, W. P. (1973). In vitro cultivation of human tumors: establishment of cell lines derived from a series of solid tumors. J Natl Cancer Inst, 51(5), 1417–1423. doi: 10.1093/jnci/51.5.1417

Graf, L., Dick, A., Sendker, F., Barth, E., Marz, M., Daumke, O., & Kochs, G. (2018). Effects of allelic variations in the human myxovirus resistance protein A on its antiviral activity. J Biol Chem, 293(9), 3056–3072. doi: 10.1074/jbc.M117.812784

Graham, F. L., Smiley, J., Russell, W. C., & Nairn, R. (1977). Characteristics of a human cell line transformed by DNA from human adenovirus type 5. J Gen Virol, 36(1), 59–74. doi: 10.1099/0022-1317-36-1-59

Hernandez, N., Melki, I., Jing, H., Habib, T., Huang, S. S. Y., Danielson, J., … Casanova, J. L. (2018). Life-threatening influenza pneumonitis in a child with inherited IRF9 deficiency. J Exp Med, 215(10), 2567–2585. doi: 10.1084/jem.20180628

Herrera-Ramos, E., Lopez-Rodriguez, M., Ruiz-Hernandez, J. J., Horcajada, J. P., Borderias, L., Lerma, E., … Rodriguez-Gallego, C. (2014). Surfactant protein A genetic variants associate with severe respiratory insufficiency in pandemic influenza A virus infection. Crit Care, 18(3), R127. doi: 10.1186/cc13934

Hidaka, F., Matsuo, S., Muta, T., Takeshige, K., Mizukami, T., & Nunoi, H. (2006). A missense mutation of the Toll-like receptor 3 gene in a patient with influenza-associated encephalopathy. Clin Immunol, 119(2), 188–194. doi: 10.1016/j.clim.2006.01.005

Hong, R., Macfarlan, T., Kutney, S. N., Seo, S.-b., Mukai, Y., Yelin, F., … Chakravarti, D. (2004). The Identification of Phosphorylation Sites of pp32 and Biochemical Purification of a Cellular pp32-kinase. Biochemistry, 43(31), 10157–10165. doi: 10.1021/bi0493968

Huyton, T., & Wolberger, C. (2007). The crystal structure of the tumor suppressor protein pp32 (Anp32a): structural insights into Anp32 family of proteins. Protein Sci, 16(7), 1308–1315. doi: 10.1110/ps.072803507

Kaltenegger, E., & Ober, D. (2015). Paralogue Interference Affects the Dynamics after Gene Duplication. Trends Plant Sci, 20(12), 814–821. doi: 10.1016/j.tplants.2015.10.003

Karczewski, K. J., Francioli, L. C., Tiao, G., Cummings, B. B., Alföldi, J., Wang, Q., … MacArthur, D. G. (2019). Variation across 141,456 human exomes and genomes reveals the spectrum of loss-of-function intolerance across human protein-coding genes. bioRxiv, 531210. doi: 10.1101/531210

Karlsson, E. K., Kwiatkowski, D. P., & Sabeti, P. C. (2014). Natural selection and infectious disease in human populations. Nature Reviews Genetics, 15(6), 379–393. doi: 10.1038/nrg3734

Kleiner, R. E., Hang, L. E., Molloy, K. R., Chait, B. T., & Kapoor, T. M. (2018). A Chemical Proteomics Approach to Reveal Direct Protein-Protein Interactions in Living Cells. Cell Chem Biol, 25(1), 110-120.e113. doi: 10.1016/j.chembiol.2017.10.001

Kondoh, T., Letko, M., Munster, V. J., Manzoor, R., Maruyama, J., Furuyama, W., … Takada, A. (2018). Single-Nucleotide Polymorphisms in Human NPC1 Influence Filovirus Entry Into Cells. J Infect Dis, 218(suppl_5), S397–s402. doi: 10.1093/infdis/jiy248

Lee, N., Cao, B., Ke, C., Lu, H., Hu, Y., Tam, C. H. T., … Chan, P. K. S. (2017). IFITM3, TLR3, and CD55 Gene SNPs and Cumulative Genetic Risks for Severe Outcomes in Chinese Patients With H7N9/H1N1pdm09 Influenza. J Infect Dis, 216(1), 97–104. doi: 10.1093/infdis/jix235

Liang, L., Jiang, L., Li, J., Zhao, Q., Wang, J., He, X., … Li, C. (2019). Low Polymerase Activity Attributed to PA Drives the Acquisition of the PB2 E627K Mutation of H7N9 Avian Influenza Virus in Mammals. mBio, 10(3). doi: 10.1128/mBio.01162-19

Lim, H. K., Huang, S. X. L., Chen, J., Kerner, G., Gilliaux, O., Bastard, P., … Zhang, S. Y. (2019). Severe influenza pneumonitis in children with inherited TLR3 deficiency. J Exp Med, 216(9), 2038–2056. doi: 10.1084/jem.20181621

Lin, Y. C., Boone, M., Meuris, L., Lemmens, I., Van Roy, N., Soete, A., … Callewaert, N. (2014). Genome dynamics of the human embryonic kidney 293 lineage in response to cell biology manipulations. Nat Commun, 5, 4767. doi: 10.1038/ncomms5767

Lindesmith, L., Moe, C., Marionneau, S., Ruvoen, N., Jiang, X., Lindblad, L., … Baric, R. (2003). Human susceptibility and resistance to Norwalk virus infection. Nat Med, 9(5), 548–553. doi: 10.1038/nm860

Long, J. S., Giotis, E. S., Moncorge, O., Frise, R., Mistry, B., James, J., … Barclay, W. S. (2016). Species difference in ANP32A underlies influenza A virus polymerase host restriction. Nature, 529(7584), 101–104. doi: 10.1038/nature16474

Long, J. S., Idoko-Akoh, A., Mistry, B., Goldhill, D., Staller, E., Schreyer, J., … Barclay, W. (2019). Species specific differences in use of ANP32 proteins by influenza A virus. eLife, 8, e45066. doi: 10.7554/eLife.45066

Lonsdale, J., Thomas, J., Salvatore, M., Phillips, R., Lo, E., Shad, S., … Moore, H. F. (2013). The Genotype-Tissue Expression (GTEx) project. Nature Genetics, 45(6), 580–585. doi: 10.1038/ng.2653

Mills, T. C., Rautanen, A., Elliott, K. S., Parks, T., Naranbhai, V., Ieven, M. M., … Hill, A. V. (2014). IFITM3 and susceptibility to respiratory viral infections in the community. J Infect Dis, 209(7), 1028–1031. doi: 10.1093/infdis/jit468

Mistry, B., Long, J. S., Schreyer, J., Staller, E., Sanchez-David, R. Y., & Barclay, W. S. (2020). Elucidating the Interactions between Influenza Virus Polymerase and Host Factor ANP32A. J Virol, 94(3), e01353–01319. doi: 10.1128/jvi.01353-19

Nogales, A., & M, L. D. (2019). Host Single Nucleotide Polymorphisms Modulating Influenza A Virus Disease in Humans. Pathogens, 8(4). doi: 10.3390/pathogens8040168

Obri, A., Ouararhni, K., Papin, C., Diebold, M. L., Padmanabhan, K., Marek, M., … Hamiche, A. (2014). ANP32E is a histone chaperone that removes H2A.Z from chromatin. Nature, 505(7485), 648–653. doi: 10.1038/nature12922

Peacock, T. P., Sheppard, C. M., Staller, E., & Barclay, W. S. (2019). Host Determinants of Influenza RNA Synthesis. Annual Review of Virology, 6(1), 215–233. doi: 10.1146/annurev-virology-092917-043339

Peacock, T. P., Sheppard, C. M., Staller, E., Frise, R., Swann, O. C., Goldhill, D. H., … Barclay, W. S. (2020). Mammalian ANP32A and ANP32B proteins drive alternative avian influenza virus polymerase adaptations. bioRxiv, 2020.2009.2003.282384. doi: 10.1101/2020.09.03.282384

Peacock, T. P., Swann, O. C., Staller, E., Leung, P. B., Goldhill, D. H., Zhou, H., … Barclay, W. S. (2020). Swine ANP32A supports avian influenza virus polymerase. bioRxiv, 2020.2001.2024.916916. doi: 10.1101/2020.01.24.916916

Peng, Q., Liu, Y., Peng, R., Wang, M., Yang, W., Song, H., … Shi, Y. (2019). Structural insight into RNA synthesis by influenza D polymerase. Nat Microbiol, 4(10), 1750–1759. doi: 10.1038/s41564-019-0487-5

Pittman, K. J., Glover, L. C., Wang, L., & Ko, D. C. (2016). The legacy of past pandemics: common human mutations that protect against infectious disease. PLoS Pathog, 12(7).

Prabhu, S. S., Chakraborty, T. T., Kumar, N., & Banerjee, I. (2018). Association between IFITM3 rs12252 polymorphism and influenza susceptibility and severity: A meta-analysis. Gene, 674, 70–79. doi: 10.1016/j.gene.2018.06.070

Randolph, A. G., Yip, W. K., Allen, E. K., Rosenberger, C. M., Agan, A. A., Ash, S. A., … Thomas, P. G. (2017). Evaluation of IFITM3 rs12252 Association With Severe Pediatric Influenza Infection. J Infect Dis, 216(1), 14–21. doi: 10.1093/infdis/jix242

Reilly, P. T., Yu, Y., Hamiche, A., & Wang, L. (2014). Cracking the ANP32 whips: important functions, unequal requirement, and hints at disease implications. Bioessays, 36(11), 1062–1071. doi: 10.1002/bies.201400058

Saavedra, F., Rivera, C., Rivas, E., Merino, P., Garrido, D., Hernández, S., … Loyola, A. (2017). PP32 and SET/TAF-Iβ proteins regulate the acetylation of newly synthesized histone H4. Nucleic Acids Res, 45(20), 11700–11710. doi: 10.1093/nar/gkx775

Sologuren, I., Martinez-Saavedra, M. T., Sole-Violan, J., de Borges de Oliveira, E., Jr., Betancor, E., Casas, I., … Rodriguez-Gallego, C. (2018). Lethal Influenza in Two Related Adults with Inherited GATA2 Deficiency. J Clin Immunol, 38(4), 513–526. doi: 10.1007/s10875-018-0512-0

Staller, E., Sheppard, C. M., Neasham, P. J., Mistry, B., Peacock, T. P., Goldhill, D. H., … Barclay, W. S. (2019). ANP32 Proteins Are Essential for Influenza Virus Replication in Human Cells. J Virol, 93(17). doi: 10.1128/jvi.00217-19

Taliun, D., Harris, D. N., Kessler, M. D., Carlson, J., Szpiech, Z. A., Torres, R., … Abecasis, G. R. (2019). Sequencing of 53,831 diverse genomes from the NHLBI TOPMed Program. bioRxiv, 563866. doi: 10.1101/563866

Thorven, M., Grahn, A., Hedlund, K.-O., Johansson, H., Wahlfrid, C., Larson, G., & Svensson, L. (2005). A Homozygous Nonsense Mutation (428G→A) in the Human Secretor (<em>FUT2</em>) Gene Provides Resistance to Symptomatic Norovirus (GGII) Infections. J Virol, 79(24), 15351–15355. doi: 10.1128/jvi.79.24.15351-15355.2005

Tochio, N., Umehara, T., Munemasa, Y., Suzuki, T., Sato, S., Tsuda, K., … Yokoyama, S. (2010). Solution structure of histone chaperone ANP32B: interaction with core histones H3-H4 through its acidic concave domain. J Mol Biol, 401(1), 97–114. doi: 10.1016/j.jmb.2010.06.005

Wandzik, J. M., Kouba, T., & Cusack, S. (2020). Structure and Function of Influenza Polymerase. Cold Spring Harb Perspect Med. doi: 10.1101/cshperspect.a038372

Zhang, H., Zhang, Z., Wang, Y., Wang, M., Wang, X., Zhang, X., … Wang, X. (2019). Fundamental Contribution and Host Range Determination of ANP32A and ANP32B in Influenza A Virus Polymerase Activity. J Virol, 93(13). doi: 10.1128/jvi.00174-19

Zhang, Q. (2020). Human genetics of life-threatening influenza pneumonitis. Hum Genet. doi: 10.1007/s00439-019-02108-3

Zhang, Y.-H., Zhao, Y., Li, N., Peng, Y.-C., Giannoulatou, E., Jin, R.-H., … Dong, T. (2013). Interferon-induced transmembrane protein-3 genetic variant rs12252-C is associated with severe influenza in Chinese individuals. Nature Communications, 4(1), 1418. doi: 10.1038/ncomms2433

